# Unveiling a Potential Prognostic Biomarker in KRAS-Mutant Lung Adenocarcinoma Using Weighted Gene Co-Expression Network Analysis

**DOI:** 10.1101/2023.07.31.551246

**Authors:** Yasmeen Dodin

**Author notes:** **Correspondence:** Address correspondence and reprint request to Yasmeen Dodin, Cancer Control Office, King Hussein Cancer Center, Amman 11941, Jordan.

## Abstract

**Background:** Globally, lung cancer is the leading cause of cancer-related deaths, primarily non-small cell lung cancer (NSCLC). Kirsten Rat Sarcoma Oncogene Homolog (KRAS) mutations are common in NSCLC and linked to a poor prognosis. Covalent inhibitors targeting KRAS-G12C mutation have improved treatment for some patients, but most KRAS-mutant lung adenocarcinoma (KRAS-MT LUAD) cases lack targeted therapies. More research is required to identify prognostic genes in KRAS-MT LUAD.

**Objective:** We aimed to identify hub genes within key co-expression gene network modules specifically associated with KRAS-MT LUAD. These hub genes hold the potential to serve as therapeutic targets or biomarkers, providing insights into the pathogenesis and prognosis of lung cancer.

**Methods:** We performed a comprehensive analysis on KRAS-MT LUAD using diverse data sources. This included TCGA project data for RNA-seq, clinical information, and somatic mutations, along with RNA-seq data for adjacent normal tissues. DESeq2 identified differentially expressed genes (DEGs), while weighted gene co-expression network analysis (WGCNA) revealed co-expression modules. Overlapping genes between DEGs and co-expression module with the highest significance were analyzed using gene set enrichment analysis (GSEA) and protein-protein interaction (PPI) network analysis. Hub genes were identified with the Maximal Clique Centrality (MCC) algorithm in Cytoscape. Prognostic significance was assessed through survival analysis and validated using the GSE72094 dataset from Gene Expression Omnibus (GEO) database.

**Results:** In KRAS-MT LUAD, 3,122 DEGs were found (2,131 up-regulated, 985 down-regulated). The blue module, among 25 co-expression modules from WGCNA, had the strongest correlation. 804 genes overlapped between DEGs and the blue module. Among twenty hub genes in the blue module, leucine-rich repeats containing G protein-coupled receptor 4 (LGR4) overexpression correlated with worse overall survival (OS) in KRAS-MT LUAD patients (P=0.012). The prognostic significance of LGR4 was confirmed using GSE72094, but surprisingly, the direction of the association was opposite to what was expected.

**Conclusion:** LGR4 is a potential prognostic biomarker in KRAS-MT LUAD. Contrasting associations in TCGA and GSE72094 datasets reveal the complexity of KRAS-MT LUAD. Further investigations are required to understand LGR4’s role in lung adenocarcinoma prognosis, especially in the context of KRAS mutations.

## 1. INTRODUCTION

The death rate from lung cancer is the highest among cancer related deaths worldwide, accounting for 1.8 million deaths in 2020 [1]. Non-small cell lung cancers (NSCLCs) account for approximately 85% of all cases of lung cancer. Specifically, adenocarcinomas comprise around 40% of NSCLC cases [2]. Within NSCLC, the most prevalent genetic alterations occur in the Kristen Rat Sarcoma Oncogene Homolog (KRAS) gene, which is found in 20-30% of NSCLC cases. Among these mutations, the substitution of Gly12 with Ala, Cys, Asp, or Val is most commonly observed. These alterations are strongly associated with tobacco smoking, resistance to chemotherapy and radiation treatments, and overall poor prognosis [3].

The KRAS protein, belonging to the small guanosine triphosphatase (GTPase) family, operates by transitioning between two states: the active form bound to GTP and the inactive form associated with GDP. When in the active GTP-bound state, KRAS activates several crucial cellular signaling pathways, including RAF-MEK-ERK, PI3K-AKTmTOR, and RALGDS-RA [4, 5]. Oncogenic KRAS mutations hinder the GTPase’s ability to hydrolyze GTP, leaving KRAS in a constantly active state. This disruption results in uncontrolled cell division and enhanced cell survival, contributing to the development and progression of cancer [6].

Previously, it was widely believed that KRAS, due to its high affinity for GTP and the absence of clearly defined binding pockets for allosteric inhibitors, was considered untargetable and undruggable. However, this perspective underwent a significant shift with the advent of two breakthrough covalent inhibitors, adagrasib and sotorasib, specifically designed to target the KRAS-G12C mutation. These inhibitors proved to be a game-changer in the field. Before the era of KRAS-G12C inhibitors, researchers explored various therapeutic strategies aimed at improving lung cancer outcomes. These approaches involved targeting KRAS upstream regulators and downstream effectors, hoping to find alternative ways to disrupt the aberrant signaling pathways associated with KRAS-driven cancers. However, the advent of adagrasib and sotorasib marked a significant milestone, offering a direct and specific approach to inhibit KRAS-G12C and potentially revolutionize the treatment landscape for patients with lung cancer.

Adagrasib and sotorasib have demonstrated their selective and irreversible binding to the cysteine at codon 12, specifically within the KRAS mutation. This binding effectively traps KRAS in an inactive state, bound to GDP [7, 8]. It is important to note that, like other kinase inhibitors, the response to both adagrasib and sotorasib is not long-lasting due to the development of resistance [9, 10]. It is worth mentioning that KRAS-G12C mutations are only found in approximately 40% of patients with KRAS mutations, leaving more than half of patients with KRAS mutations without a targeted treatment option [11].

The rapid progress in genomics and high-throughput technologies has significantly expedited the advancement of targeted therapies, including KRAS-G12C inhibitors. These inhibitors have brought about a revolutionary shift in the treatment approach and overall outcomes for certain patients with lung cancer. However, it is crucial to note that the majority of lung cancer cases are diagnosed at advanced stages when surgical interventions are no longer viable, and the absence of approved targeted therapies as first-line treatments for KRAS-mutant metastatic NSCLC [12, 13] further limits the available treatment options. To address these challenges and improve patient care, further genomic research is required to gain a deeper understanding of KRAS genetic alterations and identify key prognostic genes in KRAS-mutant lung adenocarcinoma (KRAS-MT LUAD) that can serve as therapeutic targets or assist in facilitating the routine diagnosis and treatment of lung cancer (predictive biomarkers).

Weighted gene co-expression analysis (WGCNA) is a valuable scientific technique used to identify co-expressed gene modules and important biomarkers [14]. This method allows for the identification of modules containing genes that exhibit high correlations. Furthermore, it enables the exploration of the relationship between these modules and various clinical traits. In our analysis, we aimed to identify hub genes within key co-expression gene network modules specifically associated with KRAS-MT LUAD. These hub genes hold the potential to serve as therapeutic targets or biomarkers, providing insights into the pathogenesis and prognosis of lung cancer.

## 2. MATERIALS AND METHODS

Figure 1. presents a broad outline of the current analysis workflow. Further elaborations on the specific analyses can be found in the subsequent sections.

**Figure 1.**
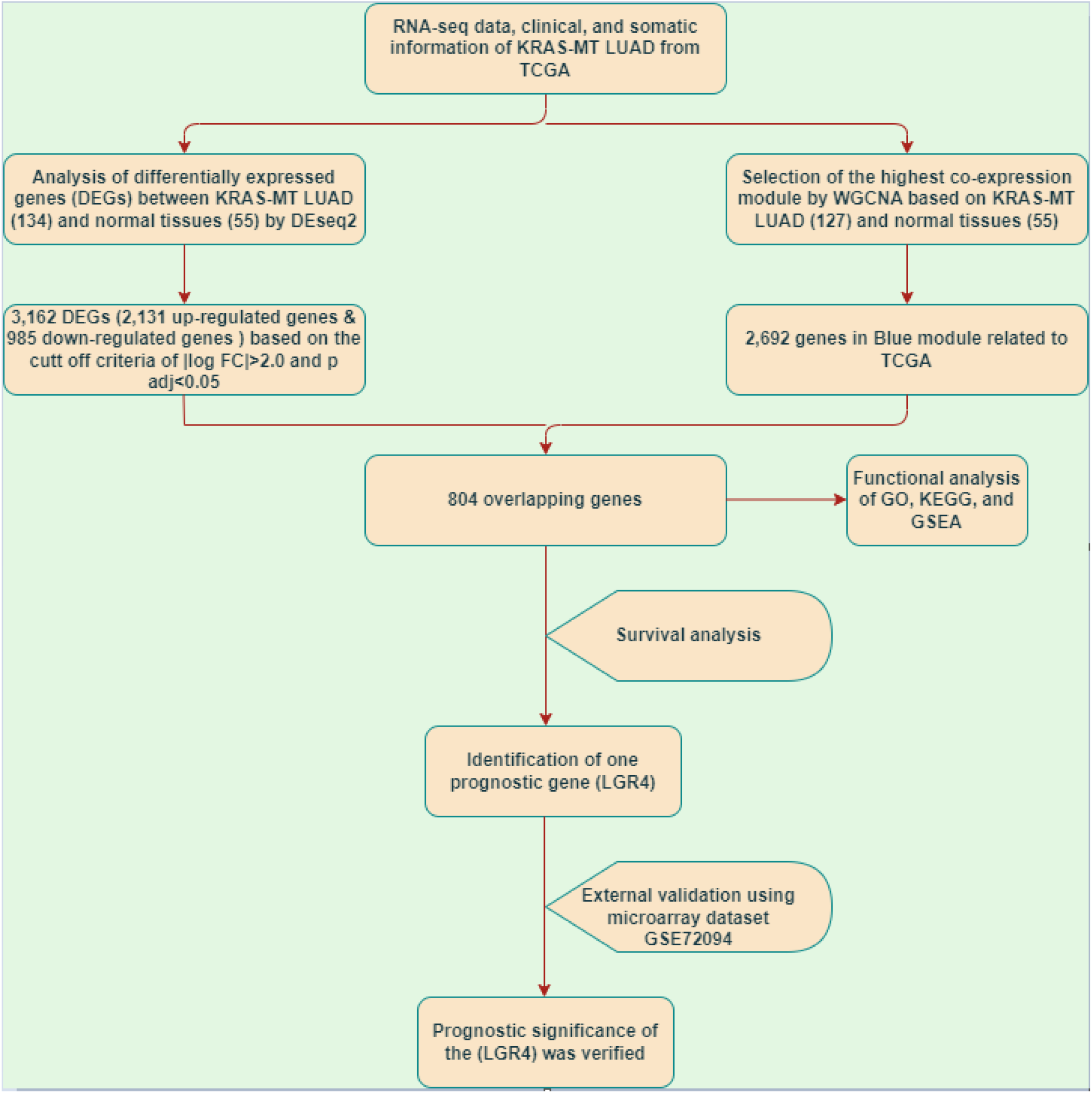

### 2.1 RNA-seq data, clinical, and somatic mutation information

The KRAS-MT LUAD RNA-seq data, along with corresponding clinicopathological features and somatic mutation information, were retrieved from the Cancer Genome Atlas Lung Adenocarcinoma (TCGA-LUAD) project [15]. The retrieval process utilized the TCGAbiolinks R/Bioconductor package [16], as well as the Maftools Bioconductor package [17]. For the TCGAbiolinks package, the retrieval conditions were similar to what specified in our previous work [18], with the exception of *sample.type* (Primary Tumor and Solid Tissue Normal were used instead of Primary Tumor) and ajcc_pathologic_stage (all stages were included instead of advanced stages). A total of 134 KRAS-MT LUAD samples and 55 normal samples were retrieved for subsequent analyses.

### 2.2 Differentially expressed genes (DEGs) screening

The DESeq2 package in R was employed to identify the DEGs between KRAS-MT LUAD and normal tissues, using raw read counts [19]. To mitigate potential bias arising from genes with low expression counts, genes with values below ten were excluded from the analysis. The criteria for identifying DEGs included an absolute log2 fold change (|log2 FC|) of ≥2 and an adjusted P value of ≤0.05. The findings were visualized through an MA plot (created using the plotMA function in DESeq2), a volcano plot, and a heatmap (generated using the ggplot2 package in R) [20].

### 2.3 Identification of KRAS-MT LUAD related modules with WGCNA

In this study, the TCGA-LUAD gene expression data, consisting of KRAS-MT samples and normal tissue samples, were utilized to construct a weighted co-expression network. The construction of the network was performed using the “WGCNA” package in R [14]. To ensure the reliability of the network structure, seven outlier samples from the KRAS-MT LUAD dataset were excluded from the analysis (Figure 2). To achieve a scale-free topology and enhance the correlation adjacency matrix, a soft-thresholding power of β=8 (R2=0.89) was chosen. The expression matrix was transformed into an adjacency matrix, employing Pearson correlation coefficients to estimate the coexpression relationships between gene pairs. Subsequently, the adjacency matrix was converted into a topological overlap matrix (TOM). The DynamicTreeCut algorithm was applied, using TOM dissimilarity (diss TOM), to obtain the initial set of modules. Correlated modules (with a correlation coefficient of r ≥ 0.75) were merged to form similar modules. Module eigengenes (ME), representing the first principal components, were calculated and considered representative of the respective module’s genes. The Spearman’s correlation test was employed to assess the correlation between each eigengene and the clinical trait of interest (KRAS-MT LUAD). Further analysis focused on the genes within the most relevant KRAS-MT LUAD module.

**Figure 2.**
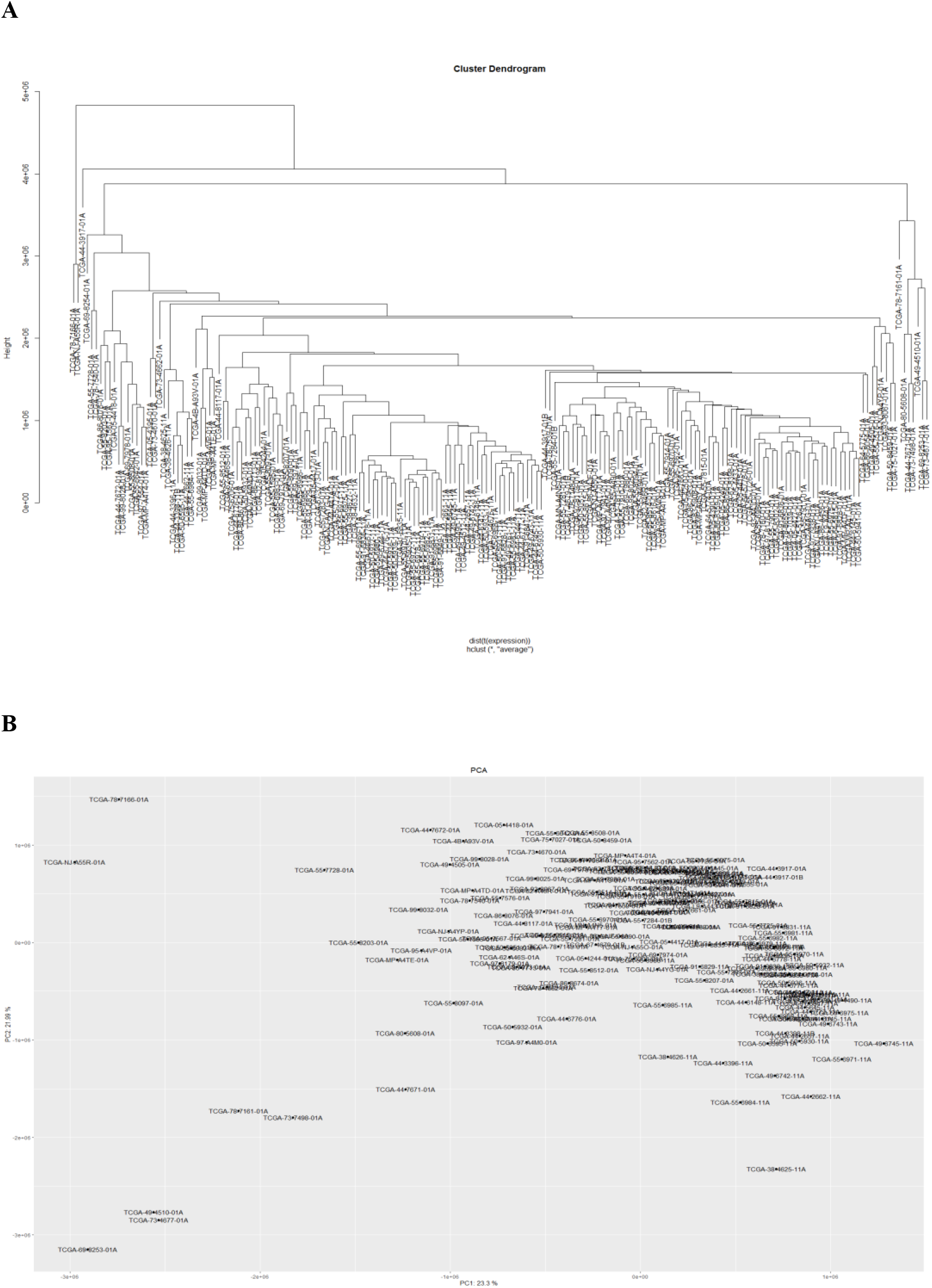
Hierarchical clustering method (A) and principal component analysis method (B) for detecting outlier samples.

### 2.4 Intersection of DEGs and WGCNA

In the subsequent steps, we focused on utilizing the most pertinent module and genes associated with KRAS-MT LUAD. These selections were made by employing WGCNA and identifying the genes that overlapped with the DEGs.

### 2.5 Application of gene set enrichment analysis (GSEA), and construction of protein-protein Interactions (PPI)

The overlapping/intersecting genes obtained from the analysis were subjected to Gene Ontology (GO) enrichment analysis, which included molecular function (MF), cellular component (CC), and biological process (BP), as well as Kyoto Encyclopedia of Genes and Genomes (KEGG) enrichment analysis. This analysis was conducted using the “clusterProfiler” package in R [21] using a minimum gene set of twenty and a P value of 0.05. Graphical representations of the significantly enriched BP, CC, MF, and KEGG pathways were generated using the ggplot2 package in R, focusing on the top five results [20]. The intersecting genes obtained from the analysis were subjected to further analysis using the STRING (Search Tool for the Retrieval of Interacting Genes) online tool [22]. This analysis aimed to construct a PPI network with a cut-off criterion of a combined score ≥ 0.9, indicating high confidence in the interactions. The resulting PPI network was visualized using Cytoscape software (v3.9.1) [23].To identify the most important genes within the co-expression network, the Maximal Clique Centrality (MCC) algorithm provided by the CytoHubba plugin in Cytoscape was applied [24]. This analysis allowed for the identification of the top twenty hub genes in the co-expression (PPI) network, which play significant roles in the network based on their connectivity and centrality.

### 2.6 Hub genes Survival analysis

To investigate the potential association of these hub genes with Overall Survival (OS) in KRAS-MT LUAD patients, the expression levels of each hub gene were categorized into two groups: patients with expression levels greater than or equal to the median, and patients with expression levels smaller than the median. To assess the survival differences between the high and low expression groups, the R packages “survival” (v3.5-5) and “survminer” (v0.4.9) were utilized. Kaplan-Meier (KM) estimates were used to analyze the survival probabilities, and the Log-rank (LR) test was employed to determine the statistical significance. A P value less than 0.05 was considered indicative of a significant difference in survival between the high and low expression groups.

### 2.7 External validation of survival-related hub genes

For external validation, we used a microarray dataset (GSE72094) to verify survival significance of the survival related hub gene. The GSE72094 dataset enrolled a total of 442 patients, all of whom had comprehensive mRNA expression data and underwent Sanger sequencing analysis to investigate the presence of EGFR, KRAS, TP53, and STK11 genes [25]. For the survival analysis we only included KRAS positive samples (N=139).

## 3. RESULTS

### 3.1 TCGA gene expression data

A comprehensive analysis was conducted on a total of 182 samples, consisting of 127 KRAS-MT LUAD samples and 55 adjacent normal tissues, to identify the DEGs. Among the screened genes, a total of 3,162 DEGs met the criteria for selection, with 2,131 genes showing up-regulation and 985 genes showing downregulation. The DEGs were visually represented through the MA plot (Figure 3A), volcano plot (Figure 3B), and heatmap (Figure 3C).

**Figure 3.**
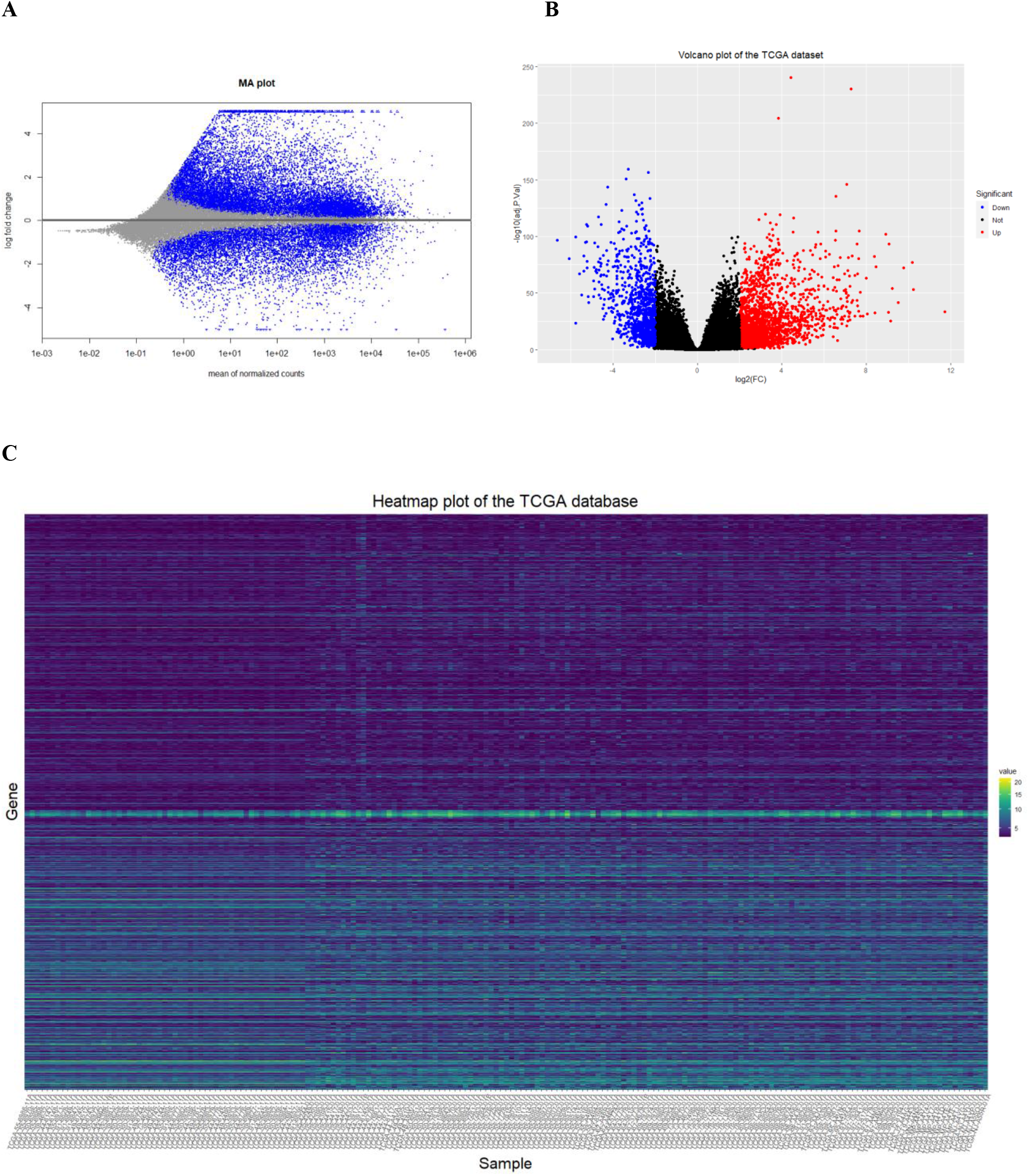
Differentially expressed genes (DEGs) in KRAS-MT LUAD. MA plot (A), volcano plot (B), heatmap (C).

### 3.2 Weighted Co-expression network and their key modules

By employing a soft-thresholding power β value of 8, the fit index curve for the scale-free topology flattened out at 0.89 (Figure 4). Subsequently, a total of 25 co-expression modules were identified within the constructed weighted gene co-expression network of KRAS-MT LUAD. These modules were obtained through the process of merging correlated modules using average linkage clustering (Figure 5).

**Figure 4.**
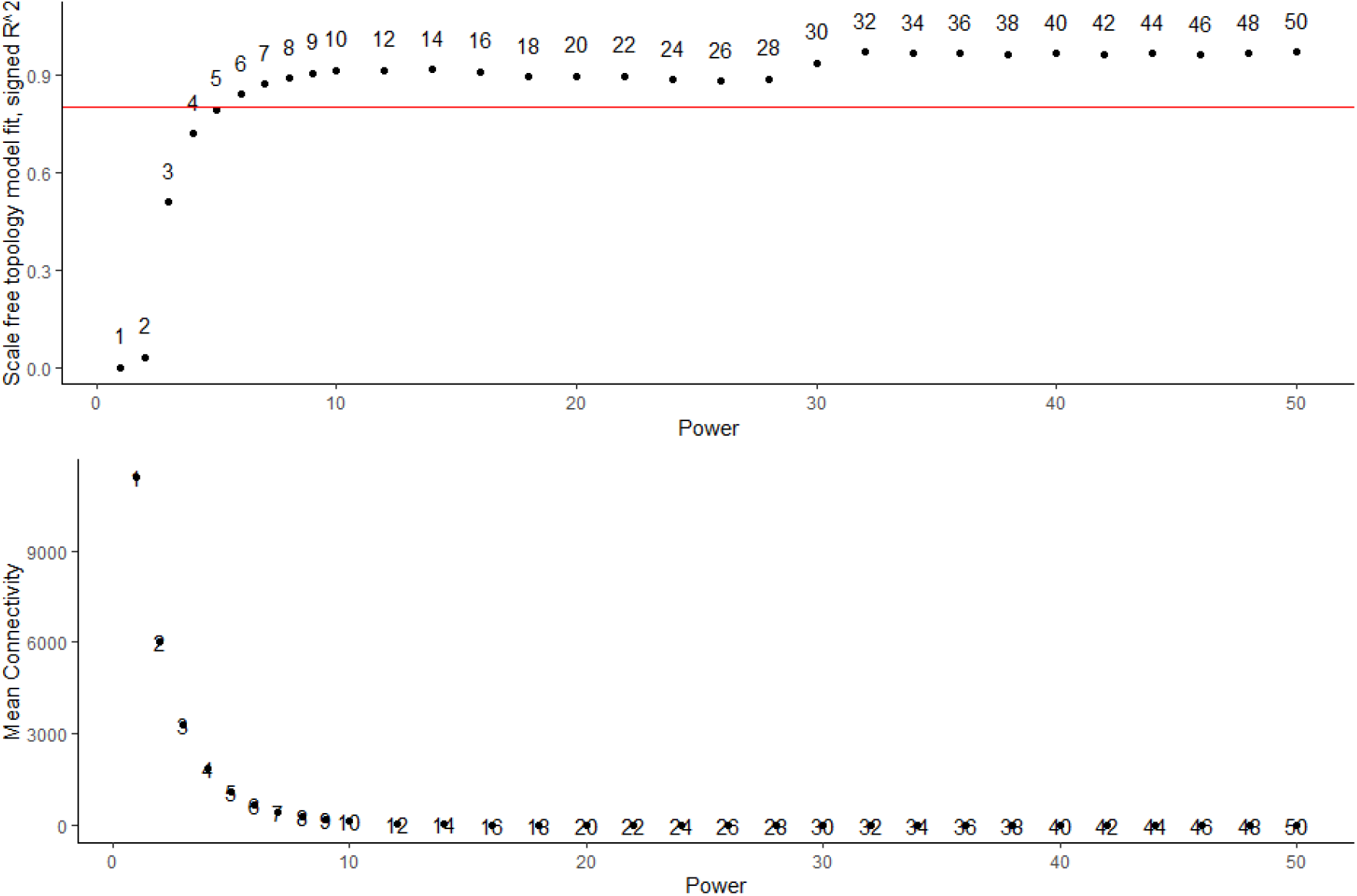
Analysis of the scale-free fit index and the mean connectivity of various soft-thresholding powers.

**Figure 5.**
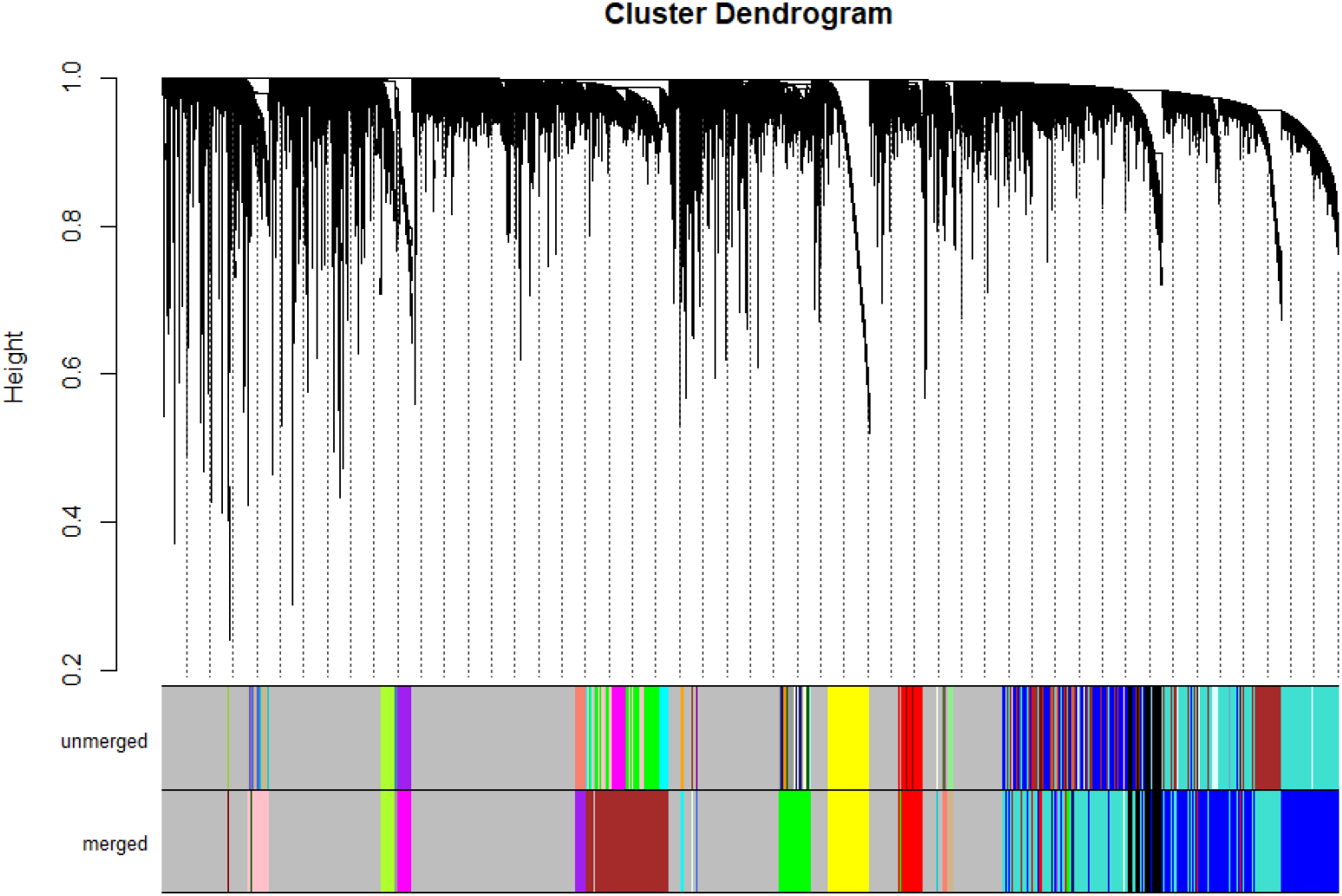
Hierarchical clustering dendograms of identified co-expressed genes in KRAS-MT LUAD.

The constructed weighted gene co-expression network in KRAS-MT LUAD comprised various modules, each represented by a distinct color such as black, light yellow, royal blue, yellow, blue, midnight blue, salmon, tan, magenta, green yellow, grey 60, dark green, pink, light green, brown, purple, light cyan, dark turquoise, green, red, turquoise, dark red, cyan, dark grey, and grey, including 21-11,221 genes. The grey module specifically consisted of genes that did not belong to any other module. Through correlation analysis between module eigengenes (ME) and KRAS-MT LUAD, it was observed that sixteen modules exhibited an association with KRAS-MT LUAD (Figure 6).

**Figure 6.**
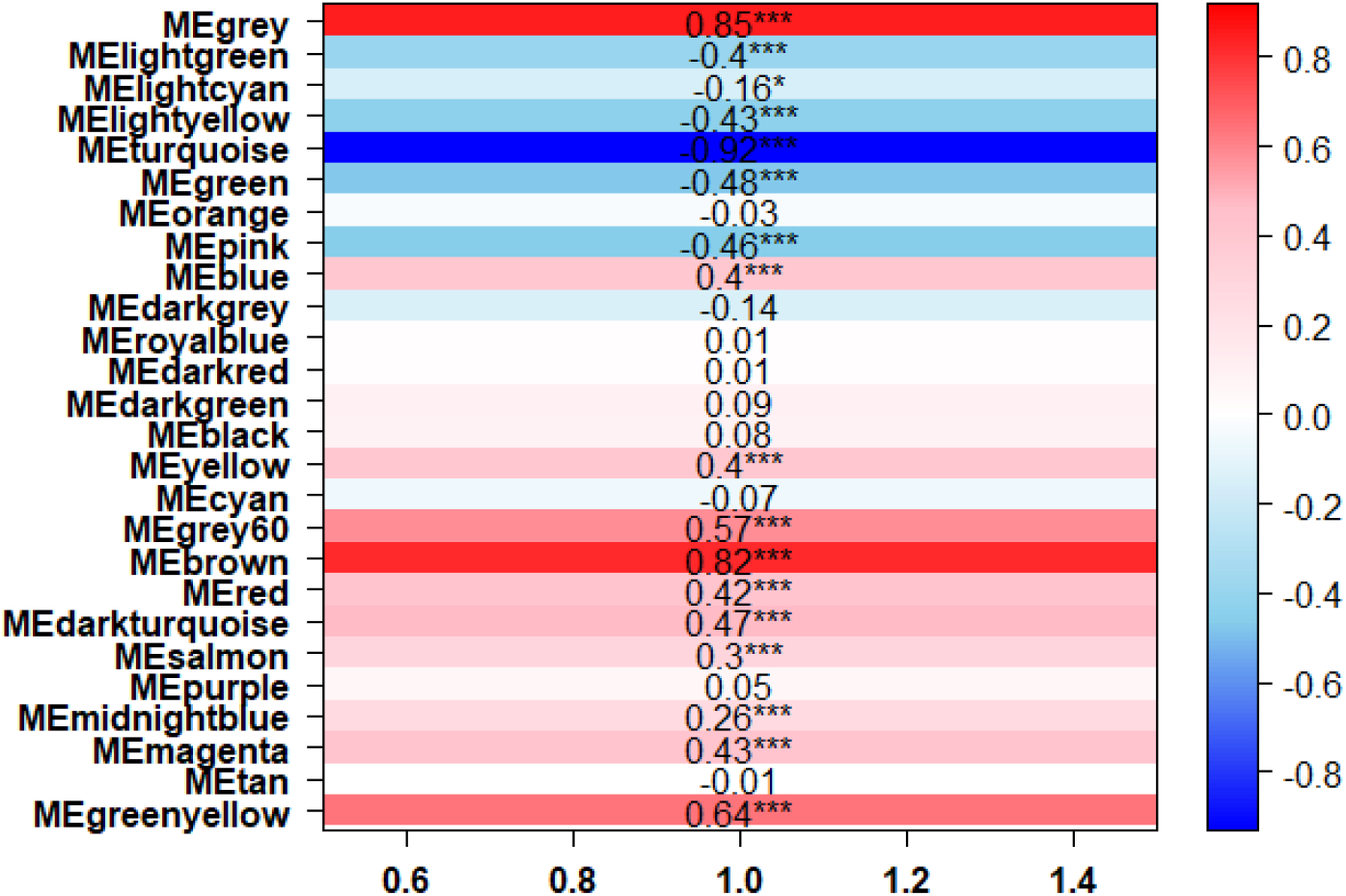
Heatmap of the correlation between Eigengene and the clinical treat of KRAS-MT LUAD.

Among the identified modules in KRAS-MT LUAD, the blue module stood out as the most significant and representative. It consisted of 2,692 genes, with a correlation (R2 = 0.40) and statistical significance (P = 1.227674e-85). Due to its prominence, the blue module was chosen for further analysis.

### 3.3 Intersection of DEGs and WGCNA

A total of 804 genes within the blue module overlapped with the set of DEGs.

### 3.4 Application of GSEA

Functional enrichment analyses using GO and KEGG were performed on the set of 804 genes. GO enrichment analysis revealed significant enrichment in one biological processes (BP) and five cellular components (CCs). KEGG enrichment analysis identified one significantly enriched pathway. The top five enriched GO terms and KEGG pathways are displayed in Figure 7.

**Figure 7.**
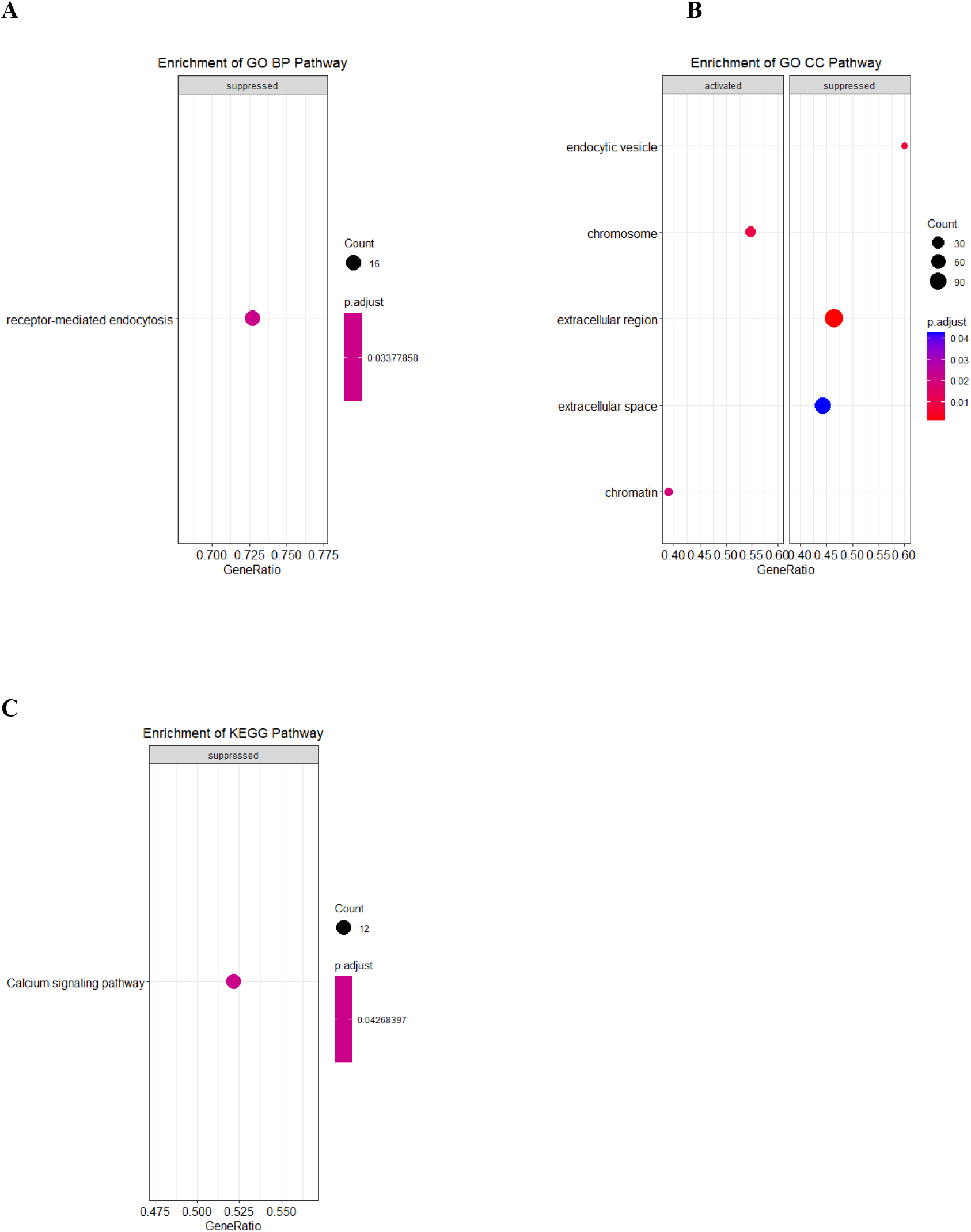
Gene set enrichment analysis (GSEA) for the DEGs intersecting with the blue module (N=804). (A) GO biological process (BP); (B) GO cellular component and (C) KEGG pathway analysis. No molecular function (MF) was significantly enriched under the specified conditions. Spot sizes represent the number of genes, while color indicates adjusted p-values.

### 3.5 Hub gene identification

With the assistance of the STRING database, a PPI network was constructed from the DEGs within the blue module. The resulting PPI network contained 804 nodes and 184 edges. To identify hub genes within the blue module, we employed the MCC algorithm through the cytoHubba plugin. The top twenty genes identified as hub genes within the blue module were ADAM Metallopeptidase With Thrombospondin Type 1 Motif 14 (ADAMTS14), ADAMTS like 3 (ADAMTSL3), Somatomedin B And Thrombospondin Type 1 Domain Containing (SBSPON), Semaphorin 5A (SEMA5A), ADAM Metallopeptidase With Thrombospondin Type 1 Motif 8 (ADAMTS8), Endothelin 1 (EDN1), Adrenoceptor Beta 2 (ADRB2), Thrombospondin Type 1 Domain Containing 1 (THSD1), Angiotensin II Receptor Type 2 (AGTR2), Cholinergic Receptor Muscarinic 1 (CHRM1), Wnt Family Member 3A (WNT3A), R-Spondin 1 (RSPO1), R-Spondin 2 (RSPO2), R-Spondin 4 (RSPO4), Leucine-rich Repeats Containing G Protein-coupled Receptor 4 (LGR4), ADAM Metallopeptidase With Thrombospondin Type 1 Motif 1 (ADAMTS1), Fibroblast Growth Factor Receptor 2 (FGFR2), Receptor Activity Modifying Protein 2 (RAMP2), Receptor Activity Modifying Protein 3 (RAMP3), and Calcitonin Receptor Like Receptor (CALCRL). These hub genes are depicted in Figure 8.

**Figure 8.**
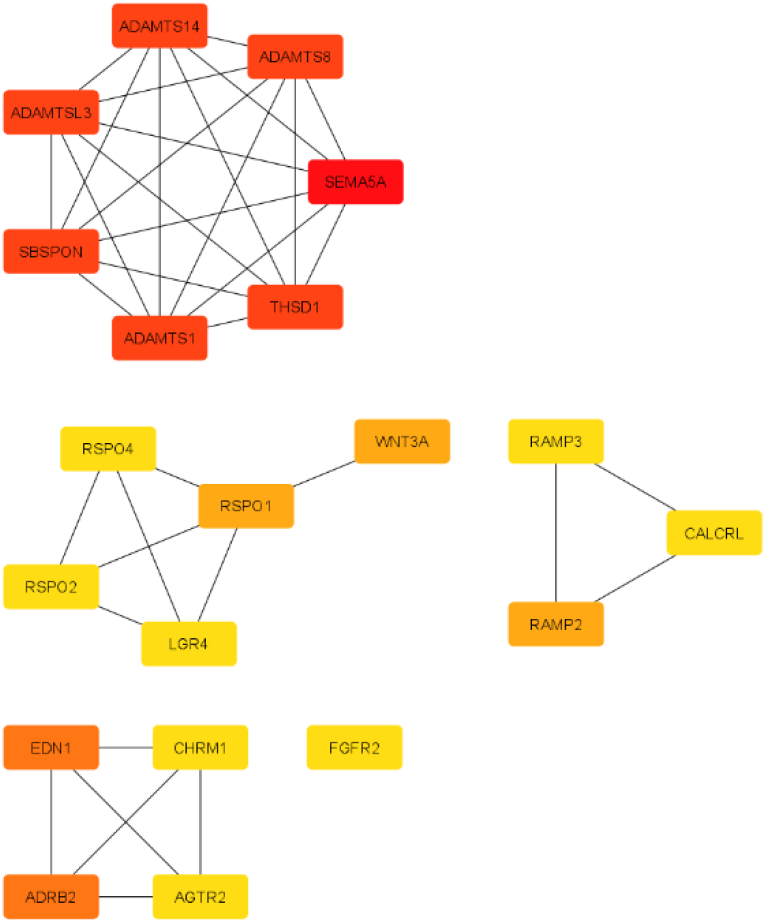
PPI analysis and identification of hub genes involved in the DEGs intersecting with the blue module using STRING database and cytoHubba plug-in Cytoscape.

### 3.6 Hub genes Survival analysis

The analysis of the twenty hub genes within the blue module revealed a significant correlation between the increased expression of one gene, LGR4, and a lower OS rate in KRAS-MT LUAD patients (P=0.012). This finding is illustrated in Figure 9.

**Figure 9.**
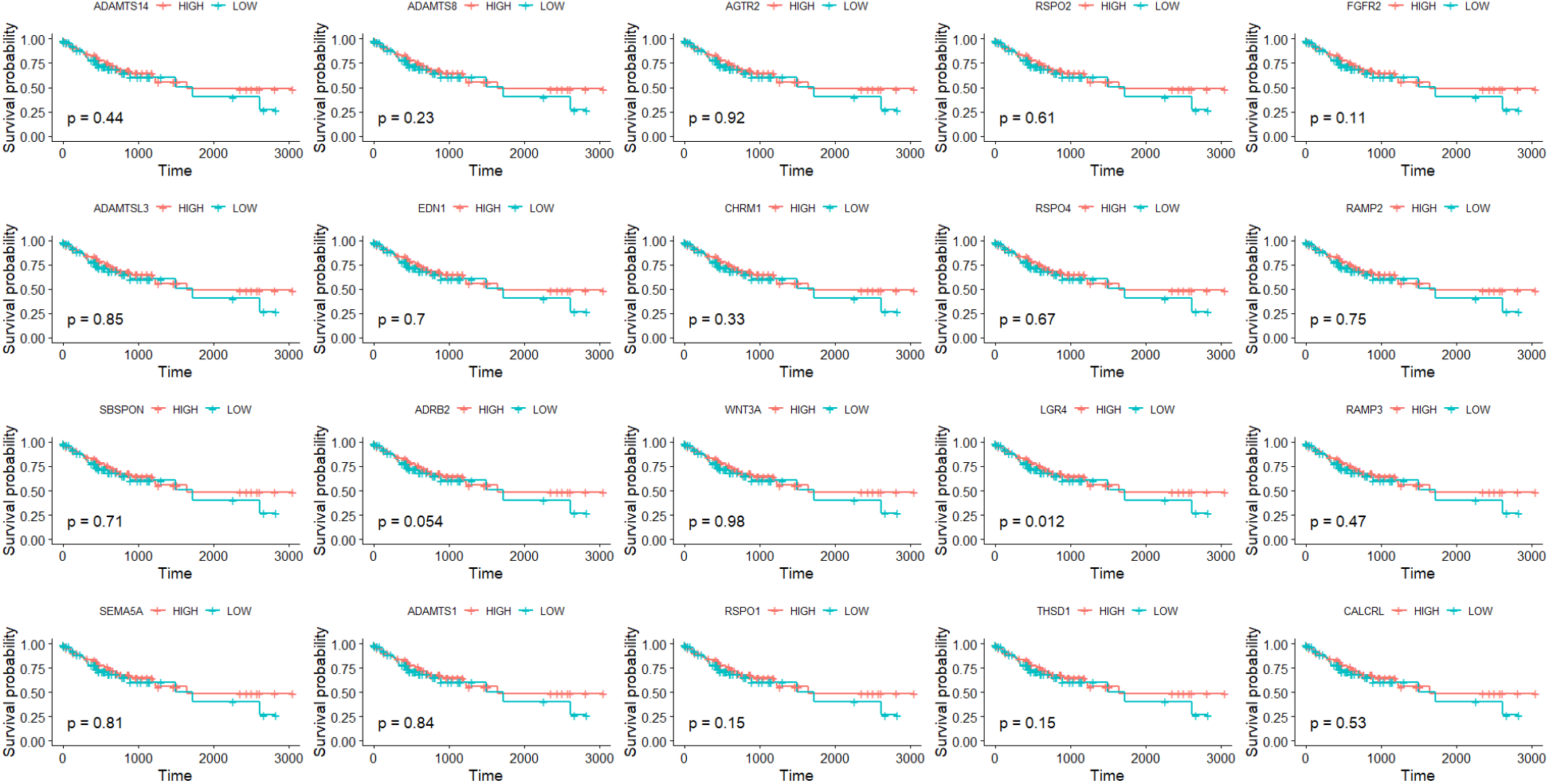
Kaplan-Meier survival of hub genes. The Patients were categorized into high-level (red) and low-level (green) groups based on the gene’s median expression.

### 3.7 External validation of the survival-related hub genes

In our study, we conducted survival analysis using RNA-seq data from KRAS-MT LUAD tissues. Our analysis identified a specific up-regulated DEG that overlapped with a tumor-related module known as the blue module. Remarkably, this gene exhibited a strong correlation with poor survival outcomes in LUAD patients. To validate the prognostic significance of this survival-related hub gene (LGR4), we sought external validation using a separate microarray dataset, GSE72094. The validation process involved performing survival analysis in a manner consistent with our previous approach using TCGA data. The results obtained from this external dataset are presented in Figure 10, further supporting the survival significance of LGR4 as observed in our initial analysis, but with the opposite direction (low expression levels of LGR4 were associated with poor survival).

**Figure 10.**
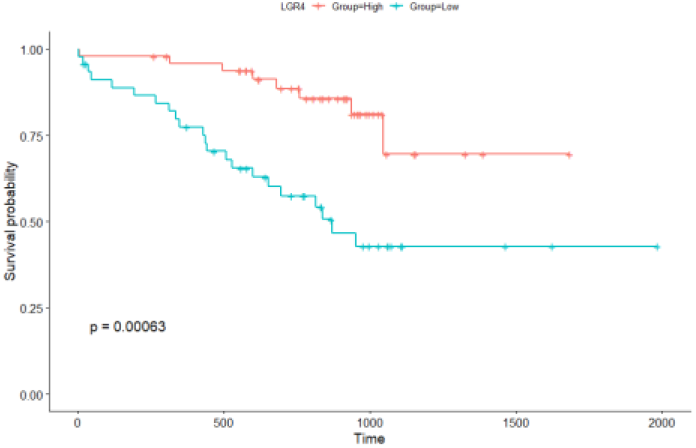
Kaplan-Meier survival of survival-related hub genes. The Patients were categorized into high-level (red) and low-level (green) groups based on the gene’s median expression.

## 4. DISCUSSION

In this study, we aimed to identify potential prognostic biomarkers in KRAS-MT LUAD using WGCNA. Our analysis revealed the blue module as the most significant and representative module associated with in KRAS-MT LUAD, consisting of 2,692 genes. We further explored the intersection of DEGs caused by KRAS mutation and the blue module, leading to the identification of 804 overlapping genes. Functional enrichment analysis using GO and KEGG pathways provided insights into the potential biological functions of these intersecting genes. In our study, we focused on identifying hub genes within the DEGs located in the blue module of in KRAS-MT LUAD. Hub genes are highly connected genes within a network and are often critical in maintaining network integrity and functionality. The top twenty hub genes identified in our analysis may hold potential as therapeutic targets or biomarkers in in KRAS-MT LUAD. To evaluate the prognostic significance of the identified hub genes, we performed survival analysis and categorized patients into high and low expression groups based on the median expression levels of each hub gene. The survival analysis indicated a potential correlation between the expression of LGR4 and the OS of patients with in KRAS-MT LUAD, where elevated levels of LGR4 were associated with poor survival outcomes. LGR4, also referred to as protein-coupled receptor 87 (GPR87), belongs to the group B of the LGR family, which includes LGR4, LGR5, and LGR6 receptors. These transmembrane receptors are members of the superfamily of G protein-coupled receptors (GPCRs), play significant roles in developmental processes, and have implications in various cancer types [26].

To validate our findings, we performed an external validation using a microarray dataset (GSE72094) of KRAS-MT LUAD. This validation aimed to assess the survival significance of the identified survival-related hub gene. Our analysis of the microarray dataset GSE72094 has uncovered a perplexing discrepancy that challenges the prevailing understanding of LGR4’s role in LUAD prognosis. Few studies have focused on the prognostic value of LGR4, while the majority of these studies and analyses [27-30], including our own TCGA analysis, have consistently associated higher expression levels of LGR4 with worse OS in LUAD patients, our validation results based on the microarray dataset revealed a strikingly contrasting pattern. Surprisingly, we observed that lower expression levels of LGR were correlated with poor OS, in direct opposition to the established consensus. Notably, an earlier study by Rao et al. [31] also presented similar findings, providing further support for our validation results. They demonstrated that in KRAS lung cancer, the gene expression patterns associated with an active receptor activator of nuclear factor-kB (RANK)/receptor activator of nuclear factor-κB ligand (RANKL) system are linked to worse OS. LGR4, which acts as a membrane-bound negative regulator by countering RANK activation, is a receptor for RANKL [32]. This observation could provide an explanation for Rao et al.’s discovery that lower mRNA expression of LGR4 is linked to an unfavorable outcome.

The existence of such divergent findings raises intriguing questions about the underlying mechanisms driving LGR4’s impact on LUAD prognosis. One possible explanation could be the inherent heterogeneity within LUAD, as different subtypes or molecular characteristics might respond differently to LGR4 expression. Moreover, KRAS mutations occurring within the same codon may exhibit non-identical biological effects, leading to variations in progression-free survival rates, changes in tumor gene expression, and differences in drug responsiveness [33]. Additionally, variations in the sample size, patient demographics, treatment regimens, and follow-up protocols across different studies may contribute to the observed disparities. Furthermore, it is crucial to consider the complexities of LGR4’s signaling pathways and its interactions with other molecular players within the tumor microenvironment.

These contrasting outcomes highlight the need for further comprehensive investigations to elucidate the factors that determine LGR4’s role in the prognosis of LUAD as a whole, and specifically in KRAS-mutant LUAD. Future functional studies should be conducted to elucidate the biological mechanisms underlying the association between LGR4 expression and survival in KRAS-MT LUAD. This could involve *in vitro* and *in vivo* experiments, such as cell line models, animal models, and molecular assays, to investigate the impact of LGR4 on tumor growth, metastasis, and response to therapy. Moreover, the association of LGR4 with clinical variables, such as stage, tumor size, lymph node involvement, and treatment response, should be assessed. This will help determine whether LGR4 expression is an independent prognostic factor or if its association with survival is confounded by other clinical factors. By gaining a deeper understanding of these complexities, we can pave the way for personalized therapeutic approaches and refine prognostic models in the context of KRAS-MT LUAD.

## CONCLUSION

In conclusion, our study identified LGR4 as a potential prognostic biomarker in KRAS-MT LUAD using WGCNA. The differences in how LGR4 expression and survival are related in the TCGA and GSE72094 datasets emphasize the intricate nature of KRAS-MT LUAD. Further investigations are warranted to elucidate the precise role of LGR4 in lung adenocarcinoma prognosis, particularly in the context of KRAS mutations. These findings contribute to our understanding of KRAS-MT LUAD and provide insights into potential biomarkers for patient prognosis and personalized treatment strategies.

## LIST OF ABBREVIATIONS

NSCLC: Non-small cell lung cancer
KRAS: Kirsten Rat Sarcoma Oncogene Homolog
KRAS-MT LUAD: KRAS-Mutant Lung Adenocarcinoma
WGCNA: Weighted gene co-expression analysis
TCGA-LUAD: The Cancer Genome Atlas Lung Adenocarcinoma
DEGs: Differentially expressed genes
GSEA: Gene set enrichment analysis
PPI: Protein-protein interactions
GO: Gene Ontology
MF: Molecular function
CC: Cellular component
BP: Biological process
KEGG: Koyoto Encyclopedia of Genes and Genomes
STRING: Search Tool for the Retrieval of Interacting Genes
MCC: Maximal Clique Centrality
OS: Overall Survival
KM: Kaplan-Meier
ADAMTS14: ADAM Metallopeptidase With Thrombospondin Type 1 Motif 14
ADAMTSL3: ADAMTS like 3
SBSPON: Somatomedin B And Thrombospondin Type 1 Domain Containing
SEMA5A: Semaphorin 5A
ADAMTS8: ADAM Metallopeptidase With Thrombospondin Type 1 Motif 8
EDN1: Endothelin 1
ADRB2: Adrenoceptor Beta 2
THSD1: Thrombospondin Type 1 Domain Containing 1
AGTR2: Angiotensin II Receptor Type 2
CHRM1: Cholinergic Receptor Muscarinic 1
WNT3A: Wnt Family Member 3A
RSPO1: R-Spondin 1
RSPO2: R-Spondin 2
RSPO4: R-Spondin 4
LGR4: Leucine-rich Repeats Containing G Protein-coupled Receptor 4
ADAMTS1: ADAM Metallopeptidase with Thrombospondin Type 1 Motif 1
FGFR2: Fibroblast Growth Factor Receptor 2
RAMP2: Receptor Activity Modifying Protein 2
CALCRL: Calcitonin Receptor Like Receptor
GPR87: protein-coupled receptor 87
GPCRs: G protein-coupled receptors
RANK: Receptor activator of nuclear factor-kB
RANKL: Receptor activator of nuclear factor-κB ligand

## ETHICS APPROVAL AND CONSENT TO PARTICIPATE

Not applicable.

## HUMAN AND ANIMAL RIGHTS

No animals/humans were used for studies that are the basis of this research.

## CONSENT FOR PUBLICATION

Not applicable

## AVAILABILITY OF DATA AND MATERIALS

The author declares that the data and materials of this study are available within the article

## ACKNOWLEDGEMENTS

Declared none.

## REFERENCES

1. Ferlay J, Colombet M, Soerjomataram I, Parkin DM, Pineros M, Znaor A, et al. Cancer statistics for the year 2020: An overview. Int J Cancer. 2021.

2. Travis WD, Brambilla E, Burke AP, Marx A, Nicholson AG. Introduction to the 2015 World Health Organization Classification of Tumors of the Lung, Pleura, Thymus, and Heart. J Thorac Oncol. 2015; 10(9):1240–2.

3. Aviel-Ronen S, Blackhall FH, Shepherd FA, Tsao MS. K-ras mutations in non-small-cell lung carcinoma: a review. Clin Lung Cancer. 2006;8(1):30–8.

4. McCormick F. KRAS as a Therapeutic Target. Clin Cancer Res. 2015; 21(8):1797–801.

5. Bodemann BO, White MA. Ral GTPases and cancer: linchpin support of the tumorigenic platform. Nat Rev Cancer. 2008; 8(2):133–40.

6. Yang H, Liang SQ, Schmid RA, Peng RW. New Horizons in KRAS-Mutant Lung Cancer: Dawn After Darkness. Front Oncol. 2019; 9:953.

7. Fell JB, Fischer JP, Baer BR, Blake JF, Bouhana K, Briere DM, et al. Identification of the Clinical Development Candidate MRTX849, a Covalent KRAS (G12C) Inhibitor for the Treatment of Cancer. J Med Chem. 2020; 63(13):6679–93.

8. Skoulidis F, Li BT, Dy GK, Price TJ, Falchook GS, Wolf J, et al. Sotorasib for Lung Cancers with KRAS p.G12C Mutation. N Engl J Med. 2021; 384(25):2371–81.

9. Hong DS, Fakih MG, Strickler JH, Desai J, Durm GA, Shapiro GI, et al. KRAS(G12C) Inhibition with Sotorasib in Advanced Solid Tumors. N Engl J Med. 2020; 383(13):1207–17.

10. Akhave NS, Biter AB, Hong DS. Mechanisms of Resistance to KRAS (G12C)-Targeted Therapy. Cancer Discov. 2021; 11(6):1345–52.

11. Biernacka A, Tsongalis PD, Peterson JD, de Abreu FB, Black CC, Gutmann EJ, et al. The potential utility of remining results of somatic mutation testing: KRAS status in lung adenocarcinoma. Cancer Genet. 2016; 209(5):195–8.

12. NCCN. NCCN Guidelines for Patients® Non-Small Cell Lung Cancer-Metastatic. 2021. Available online: http://www.nccn.org/patients.

13. Planchard D PS, Kerr K, Novello S, Smit EF, Faivre-Finn C, Mok TS, Reck M, Van Schil PE, Hellmann MD, Peters S. ESMO Guidelines Committee. Metastatic non-small cell lung cancer: ESMO Clinical Practice Guidelines for diagnosis, treatment and follow-up. Ann Oncol. 2018 Oct 1; 29(Suppl 4):iv192-iv237. doi: 10.1093/annonc/mdy275. Erratum in: Ann Oncol. 2019 May; 30(5):863–870. PMID: 30285222.

14. Langfelder P, Horvath S. WGCNA: an R package for weighted correlation network analysis. BMC Bioinformatics. 2008; 9:559.

15. Cancer Genome Atlas Research N. Comprehensive molecular profiling of lung adenocarcinoma. Nature. 2014; 511(7511):543–50.

16. Colaprico A, Silva TC, Olsen C, Garofano L, Cava C, Garolini D, et al. TCGAbiolinks: an R/Bioconductor package for integrative analysis of TCGA data. Nucleic Acids Res. 2016; 44(8):e71.

17. Mayakonda A, Lin DC, Assenov Y, Plass C, Koeffler HP. Maftools: efficient and comprehensive analysis of somatic variants in cancer. Genome Res. 2018; 28(11):1747–56.

18. Yasmeen Dodin. Integrated Bioinformatics Approach for Disclosing Autophagy Pathway as a Therapeutic Target in Advanced KRAS Mutated/Positive Lung Adenocarcinoma. The Open Bioinformatics Journal. 2023 May; 16. doi: 10.2174/18750362-v16-2305230-2022-18.

19. Love MI, Huber W, Anders S. Moderated estimation of fold change and dispersion for RNA-seq data with DESeq2. Genome Biol. 2014; 15(12):550.

20. H W. Ggplot2: Elegant Graphics for Data Analysis. Springer-Verlag New York. ISBN 978-3-319-24277-42016.

21. Yu G, Wang LG, Han Y, He QY. clusterProfiler: an R package for comparing biological themes among gene clusters. OMICS. 2012; 16(5):284–7.

22. Szklarczyk D, Gable AL, Lyon D, Junge A, Wyder S, Huerta-Cepas J, et al. STRING v11: protein protein association networks with increased coverage, supporting functional discovery in genome-wide experimental datasets. Nucleic Acids Res. 2019; 47(D1):D607–D13.

23. Shannon P, Markiel A, Ozier O, Baliga NS, Wang JT, Ramage D, et al. Cytoscape: a software environment for integrated models of biomolecular interaction networks. Genome Res. 2003; 13(11):2498–504.

24. Chin C-H, Chen S-H, Wu H-H, Ho C-W, Ko M-T, Lin C-Y. cytoHubba: identifying hub objects and sub-networks from complex interactome. BMC Systems Biology. 2014; 8(S4).

25. Schabath MB, Welsh EA, Fulp WJ, Chen L, Teer JK, Thompson ZJ, et al. Differential association of STK11 and TP53 with KRAS mutation-associated gene expression, proliferation and immune surveillance in lung adenocarcinoma. Oncogene. 2016; 35(24):3209–16.

26. Van Loy T, Vandersmissen HP, Van Hiel MB, Poels J, Verlinden H, Badisco L, et al. Comparative genomics of leucine-rich repeats containing G protein-coupled receptors and their ligands. Gen Comp Endocrinol. 2008; 155(1):14–21.

27. Yang D LJ, Xu QY, Xia T, Xia JH. Inhibitory Effect of MiR-449b on Cancer Cell Growth and Invasion through LGR4 in Non-Small-Cell Lung Carcinoma. Curr Med Sci. 2018 Aug; 38(4):582–589. doi: 10.1007/s11596-018-1917-y. Epub 2018 Aug 20. PMID: 30128865.

28. Bai R, Zhang J, He F, Li Y, Dai P, Huang Z, et al. GPR87 promotes tumor cell invasion and mediates the immunogenomic landscape of lung adenocarcinoma. Commun Biol. 2022; 5(1):663.

29. Nii K, Tokunaga Y, Liu D, Zhang X, Nakano J, Ishikawa S, et al. Overexpression of G protein-coupled receptor 87 correlates with poorer tumor differentiation and higher tumor proliferation in non-small-cell lung cancer. Mol Clin Oncol. 2014; 2(4):539–44.

30. Kita Y, Go T, Nakashima N, Liu D, Tokunaga Y, Zhang X, et al. Inhibition of Cell-surface Molecular GPR87 With GPR87-suppressing Adenoviral Vector Disturb Tumor Proliferation in Lung Cancer Cells. Anticancer Res. 2020; 40(2):733–41.

31. Rao S, Sigl V, Wimmer RA, Novatchkova M, Jais A, Wagner G, et al. RANK rewires energy homeostasis in lung cancer cells and drives primary lung cancer. Genes Dev. 2017; 31(20):2099–112.

32. Luo J, Yang Z, Ma Y, Yue Z, Lin H, Qu G, et al. LGR4 is a receptor for RANKL and negatively regulates osteoclast differentiation and bone resorption. Nat Med. 2016; 22(5):539–46.

33. Ihle NT, Byers LA, Kim ES, Saintigny P, Lee JJ, Blumenschein GR, et al. Effect of KRAS oncogene substitutions on protein behavior: implications for signaling and clinical outcome. J Natl Cancer Inst. 2012; 104(3):228–39.

